# Identification of a Type IV CRISPR-Cas system located exclusively on *IncHI1B/IncFIB* plasmids in *Enterobacteriaceae*

**DOI:** 10.1101/536375

**Authors:** Enas Newire, Alp Aydin, Samina Juma, Virve I Enne, Adam P. Roberts

**Author notes:** Division of Biosciences, University College London, Gower Street, London, WCIE 6BT, UK. Quadram Institute, James Watson Road, Norwich, NR4 7UQ (Alp Aydin).

## Abstract

During an investigation of CRISPR carriage in clinical, multi-drug resistant*, Klebsiella pneumoniae*, a novel CRISPR-Cas system (which we have designated Type IV-B) was detected on plasmids from two *K. pneumoniae* isolates from Egypt (isolated in 2002-2003) and a single *K. pneumoniae* isolate from the UK (isolated in 2017). Sequence analysis of other genomes available in GenBank revealed that this novel Type IV-B CRISPR-Cas system was present on 28 other plasmids from various *Enterobacteriaceae* hosts and was never found on the chromosome. Type IV-B is found exclusively on *IncHI1B/IncFIB* plasmids and is associated with multiple putative transposable elements. Type IV-B has a single repeat-spacer array (CRISPR1) upstream of the *cas* loci with some spacers matching regions of conjugal transfer genes of *IncFIIK/IncFIB(K)* plasmids suggesting a role in plasmid incompatibility. Expression of the *cas* loci was confirmed in available clinical isolates by RT-PCR; indicating the system is active. To our knowledge, this is the first report describing a new subtype within Type IV CRISPR-Cas systems exclusively associated with *IncHI1B/IncFIB* plasmids.

**Importance:** Here, we report the identification of a novel subtype of Type IV CRISPR-Cas that is expressed and exclusively carried by *IncHI1B/IncFIB* plasmids in *Enterobacteriaceae*, demonstrating unique evolutionarily juxtaposed connections between CRISPR-Cas and mobile genetic elements (MGEs). Type IV-B encodes a variety of spacers showing homology to DNA from various sources, including plasmid specific spacers and is therefore thought to provide specific immunity against plasmids of other incompatible groups *(IncFIIK/IncFIB(K))*. The relationship between Type IV-B CRISPR-Cas and MGEs that surround and interrupt the system is likely to promote rearrangement and be responsible for the observed variability of this type. Finally, the Type IV-B CRISPR-Cas is likely to co-operate with other *cas* loci within the bacterial host genome during spacer acquisition.

## Introduction

Clustered Regularly Interspaced Short Palindromic Repeats (CRISPR) are widespread, bacterial adaptive, RNA-mediated, immune systems that target invading foreign DNA such as bacteriophages and conjugative plasmids^1, 2^. CRISPR functions through a three-stage process: adaptation involving acquisition of foreign DNA molecules spacers, expression and maturation of the short CRISPR RNAs (crRNAs), and the interference with a cognate invading foreign DNA molecule^3^. To date, CRISPR-Cas systems are classified into 2 classes, 5 Types (IV) and 33 subtypes^4^. The two classes are differentiated based on the effector module; class 1 utilises multi-protein effector Cas complexes, while class 2 utilise a single-protein effector (either Cas9 or Cpf1)^5,6^. All types are confirmed, or expected, to provide immunity against invading DNA, while Type III CRISPR-Cas systems can target both DNA and RNA^7,8^.

Type IV was previously called the Unknown Type, due to its rare occurrence and lack of the adaptation module, until an updated classification in 2015^7,9^. In 2017, Type IV classification was updated to show an associated repeat-spacer array for a *cas* loci that has *csf1* (*cas8-Like*), *csf2* (*cas7*), *csf3* (*cas5*), *csf4* (*dinG*) and *csf5* (*cas6-Like*) genetic arrangement, respectively^4^. Type IV is the only type to possess *csf4* (*dinG*)^4^. Type IV CRISPR–Cas systems were shown to employ crRNA-guided effector complexes in 2019^8^. It has been hypothesised that Type IV is similar to an ancestral innate immune system that gained adaptive ability by associating with a transposon-like element containing *cas1* and *cas2*^3^.

## Results

A novel CRISPR-cas family, which we have designated Type IV-B due to the presence of *dinG*, containing a repeat-spacer array, leader sequence and *cas* loci was detected on thirty-one (three clinical isolates and twenty-eight genomes in GenBank) *IncHI1B/IncFIB(Mar)* plasmids from *Enterobacteriaceae* (Figure 1A, Table S1). Type IV-B *cas* loci show homology to each other, while *csf5*(*cas6*) in Type IV-B shows 100% protein identity to Type I-E *cas6*. Type IV-B CRISPR-Cas systems could be grouped according to the presence of an IS element interrupting the *cas* loci and both groups are associated with multiple MGEs (Figure 1A). We also identified partial *cas* loci (*cas6* and *dinG*) on other *IncHI1B/IncFIB(Mar)* plasmids (Table S1). A single repeat-spacer CRISPR1^10^ array was identified upstream of all *cas* loci. The repeats have a predicted stem-loop secondary-structure likely involved in pre-crRNA processing (Figure 1B, 1C). Spacer-1 in CP018720.1 and spacer-20 in CP014776.1 correspond to *IncFIIK* conjugal transfer genes; *traN* and *traL*, respectively. The Protospacer Adjacent Motif (PAM) alignment revealed the leader-proximal repeat signature conservation (TGCC/TTAT). Finally, RT-PCR demonstrated that Type IV-B *cas* loci genes (*csf2, dinG* and *cas6*) are expressed.

**Figure 1:**
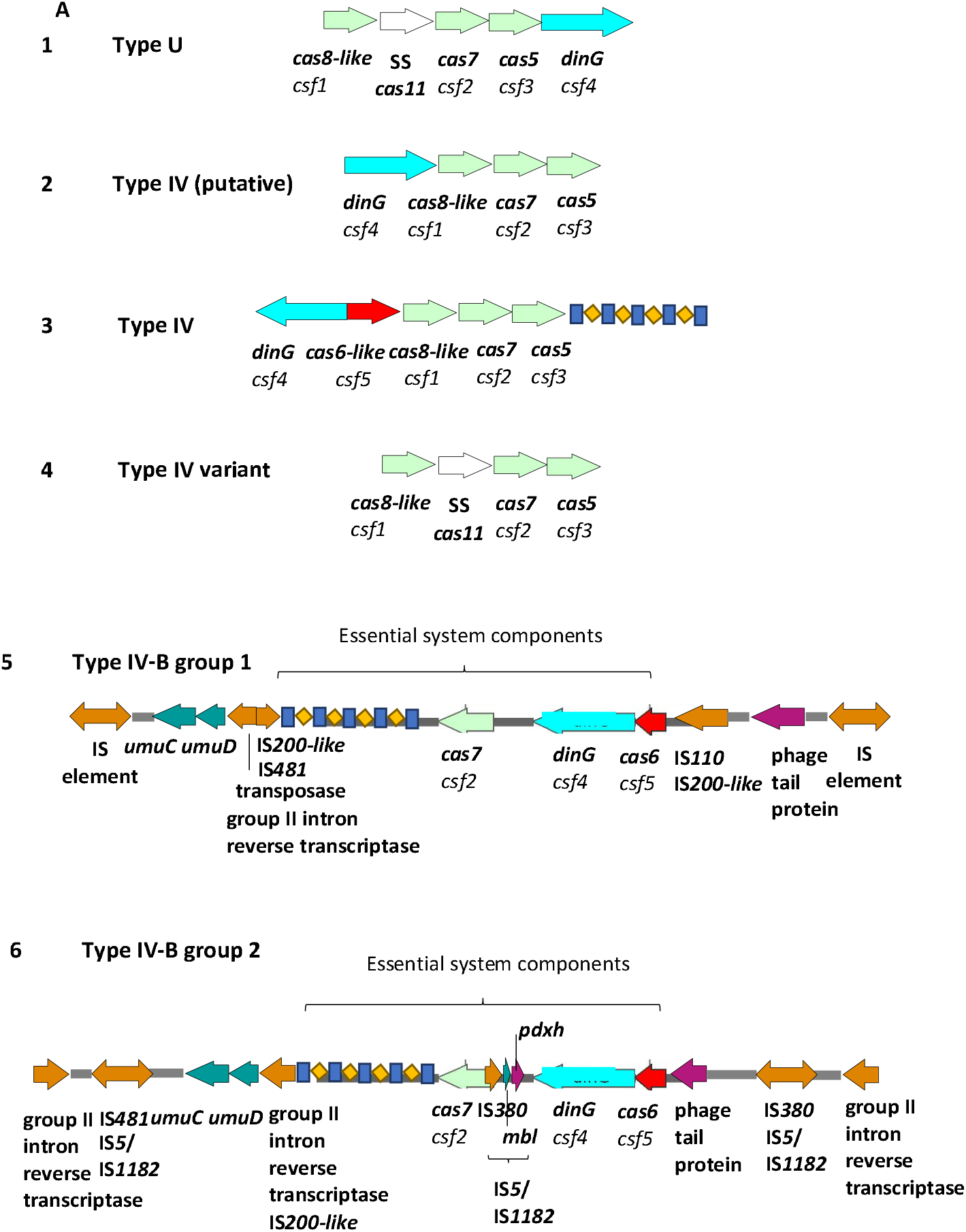
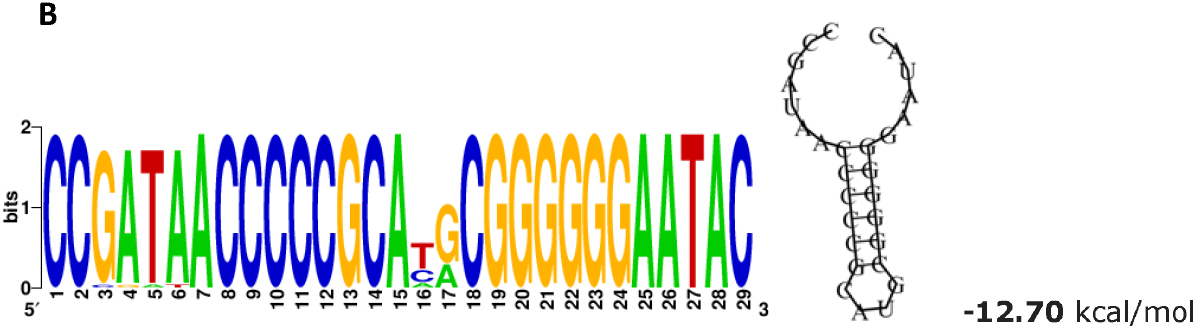
Type IV-B CRISPR-Cas system. (A) Schematic representation of Type IV CRISPR-Cas sytems and the two groups of Type IV-B described here. Panel 1 is Type U (unknown) as identified in 2013^9^. Panel 2 is Type IV (putative) as identified in *Acidithiobacillus ferroxidans* in 2015^6^. Panel 3 is Type IV and Panel 4 is Type IV variant identified in *Thioalkalivibrio sp*. K9Omix (TK90_2699-TK90_2703) and *Rhodococcus jostii* RHA1 (RHA1_ro10069-RHAl_rol0072), respectively, in 2017^14^. Panel 5 is Type IV-B group 1 and Panel 6 is Type IV-B group 2 as detected in *Enterobacteriaceae* isolates and genomes in this study. Arrows in different colours represent genes; red represents *cas6;* bright blue represents *dinG;* light green represents *cas7;* white represents *cas11;* blue-yellow pattern represents the direct repeat-spacer loci; orange represents the location of the associated MGEs occurring upstream (IS5/IS*1182*, IS*630*, IS*6*, IS*1*, IS*481* or IS*110*) and downstream (IS5/IS*1182*, IS*630*, IS*6*, IS*5-like*, IS*3000*, IS*Kra4*, IS*10L* or group II intron reverse transcriptase) of the system; green represents resistance genes; purple represents other genes associated with the system. (B) Conservation of the repeats and predicted stable stem-loop secondary structure predicted to be involved in the mechanism of pre-crRNA processing. The height of the letters in the sequence logo shows the relative frequency of their recurrence at that position. Wobbles at position 16 and 17 are within the loop of the predicted stem-loop structure and are therefore tolerated in the structural prediction shown in C; (C) The predicted secondary structure of direct repeats and the associated Minimum Free Energy (MFE) estimated in (kcal/mol) shown underneath the structure.

## Discussion

Type IV CRISPR-Cas systems are the only ones possessing *dinG*^4^, therefore we propose the CRISPR-Cas system described here be designated Type IV-B. Unlike classical Type IV, Type IV-B lacks *cas8-like* and *cas5*, however, it has *dinG* and *cas7* (involved in interference), and *cas6* (involved in expression and maturation of short crRNAs)^11–13^. Type IV-B has a variable repeat-spacer array and a conserved leader sequence. Expression of Type IV-B genes indicates system activity; likely providing immunity to incoming DNA matching the spacers^8^. The spacers demonstrated conservation and polymorphism and cluster into two main groups (Figure 2A, 2B) both matching DNA from a variety of sources. However, the adaptation module is missing, thus adding new spacers will require *cas1* and *cas2* from other CRISPR-Cas that exist within the *Enterobacteriaceae* genomes. Interestingly, some of Type IV-B spacers match conjugal transfer genes *traN* and *traL* of *IncFIIK/IncFIB(K)* plasmids, suggesting a role in plasmid incompatibility.

**Figure 2:**
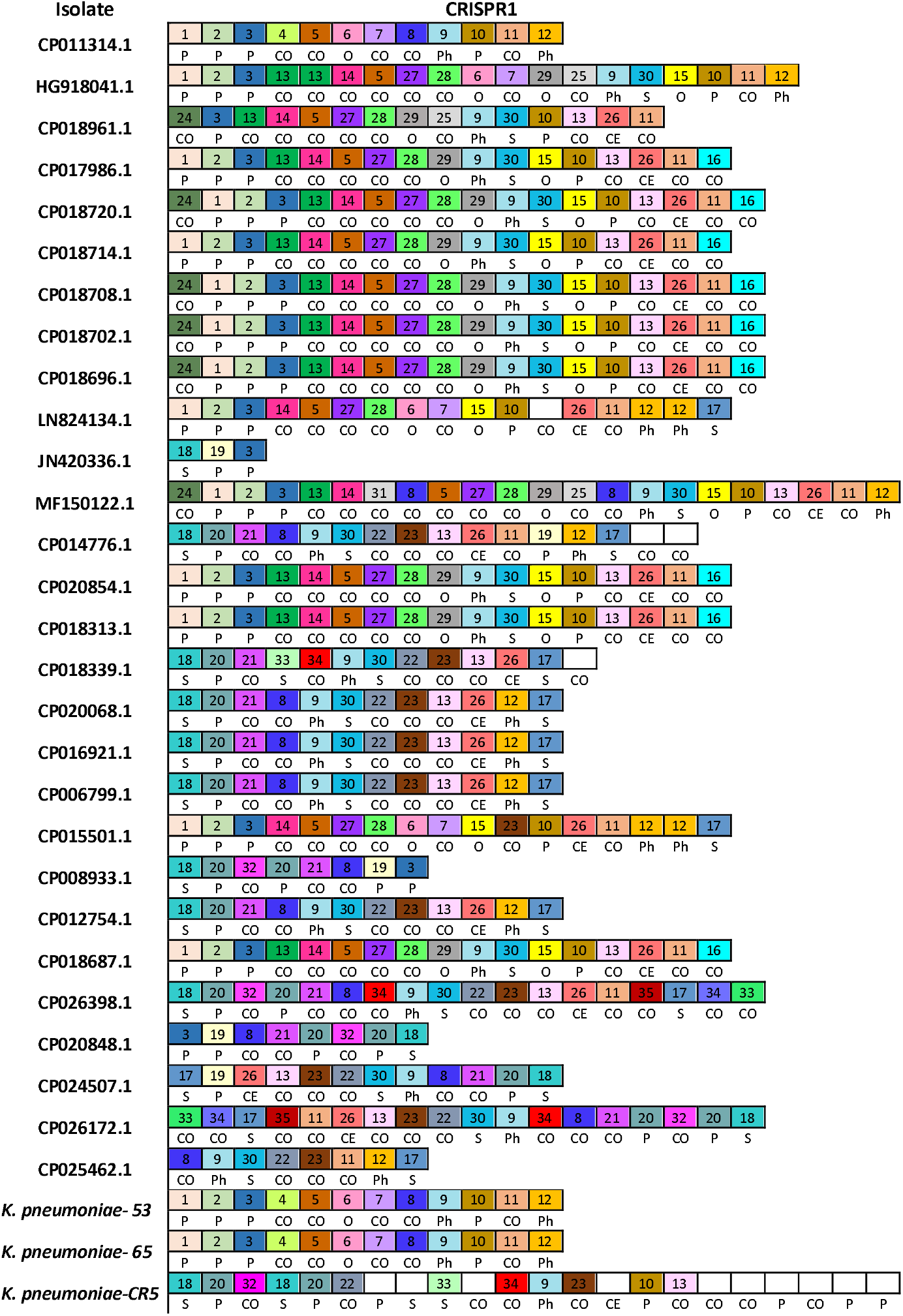
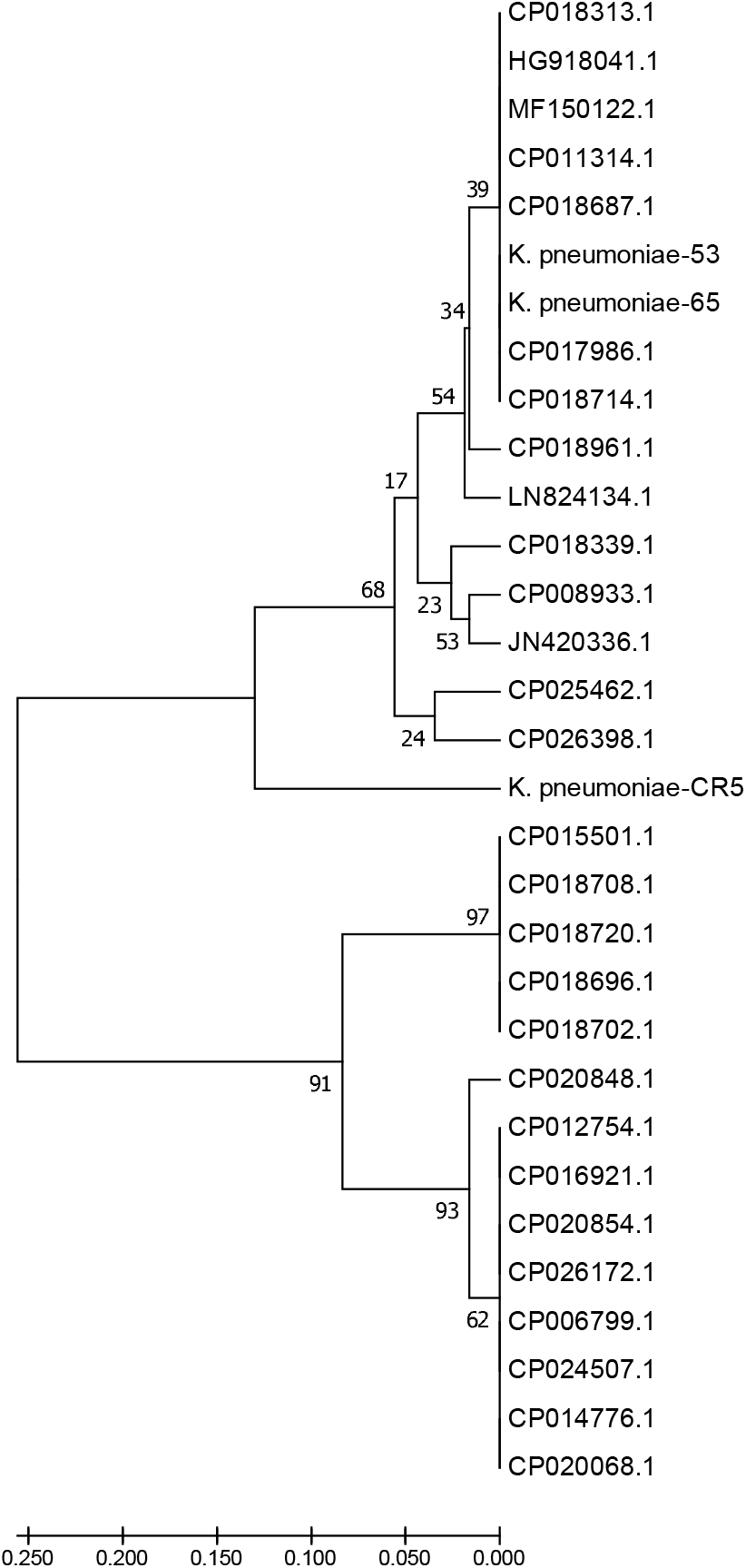
Type IV-B CRISPR spacer polymorphism and evolutionary relationships. (A) The spacers map. Only spacers are represented by boxes, and no repeats are included. Identical spacers are represented by the same number and colour, while unique spacers are represented by white colour and no number is associated with the box. Self-targeting spacers are indicated by letter (S) and show 100% identity to host DNA, plasmid-targeting spacers are indicated by letter (P), phage targeting spacers are indicated by letters (Ph), other *Enterobacteriaceae* targeting spacers (100% identity) are indicated by letter (O), cryptic spacers with similarity to other bacterial DNA are indicated by letters (CO), and those with similarity to Eukaryotic DNA are indicated by letters (CE) that are positioned underneath the relevant spacer. CE spacers showed at least 57% identity to eukaryotic DNA. CE spacers were confirmed by multiple sequences alignments. (B) The phylogenetic tree illustrating the evolutionary relationships of the Type IV-B repeat-spacer CRISPR loci. Phylogenetic UPGMA tree was constructed using the MUSCLE algorithm of MEGA7. The tree is drawn to scale, with branch lengths in the same units as those of the evolutionary distances used to infer the phylogenetic tree. The evolutionary distances were computed using the Maximum Likelihood method, [bootstrap test (1000 replicates), and the rate variation among sites was modelled with a gamma distribution (shape parameter = 1).

Type IV-B demonstrates a complex evolutionary connection with MGEs in terms of parasitism and immunity^14^. The association between Type IV-B and multiple MGEs, plus the identification of partial *cas* loci on other *IncHI1B/IncFIB(Mar)* plasmids, indicates that dynamic, MGE mediated rearrangement, of CRISPR-Cas Type IV-B is ongoing.

## Conclusion

To our knowledge, this is the first identification of a CRISPR-Cas system, which we have designated Type IV-B, exclusively associated with plasmids. The system demonstrates an evolutionary association and role for MGEs in dissemination and, additionally, the spacer analysis suggests a role in plasmid incompatibility. We propose updating the CRISPR-Cas system classification to include Type IV-B.

## Materials and Methods

### Isolate information

Clinical *K. pneumoniae-53* and *K. pneumoniae-65* were isolated from Egyptian university teaching hospitals (2002-2003), and *K. pneumoniae-CR5* from University College London Hospital in the UK (2017).

### CRISPR-Cas loci expression

RT-PCR was performed using LightCycler^®^ RNA Amplification Kit SYBR Green (Roche Diagnostics Ltd., UK). The primers were (*csf2*-fw: A A A AT GCGGTCTCAACTTCCG; *csf2*-rev:TGACGAAGAGTTCCCCGAATG), (*dinG*-fw:GAGTCTGCCGGATTGTCGTTA; *dinG-*rev:GTACCAGATAGCCCAGCGTTT) and (*cas6*-fw:AATGCGTTTCGGTTGCGTATC; cas6-rev: G AG TACG G C AG CTT CTCTCC).

### Bioinformatics analysis

DNA sequences were analysed using CRISPRFinder, CRISPRTarget and Snapgene (GSL Biotech)^12,15,16^. Multi-Locus Sequence Typing, resistance genes and plasmids were identified using MLST, ResFinder and PlasmidFinder, respectively^17^. Spacer analysis was performed by BLAST and Geneious^18^. A phylogenetic UPGMA-based tree was constructed for CRISPR using MEGA7^19, 20^. Direct repeats and PAM conservation were assessed using WebLogo, RNA secondary structure was predicted using RNAfold^21–23^.

## Transparency declarations

None to declare.

## Supplementary Table

**Table S1:**
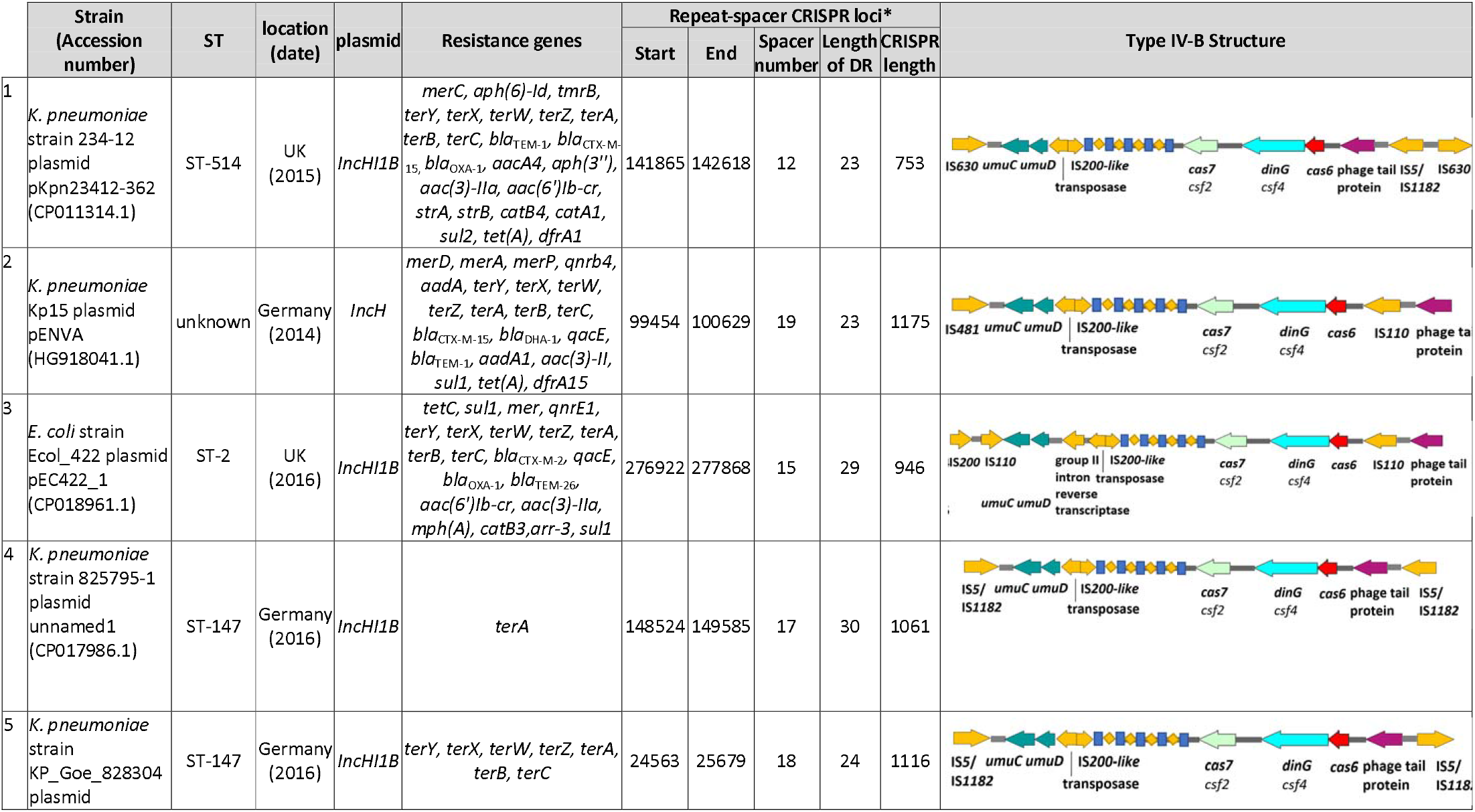
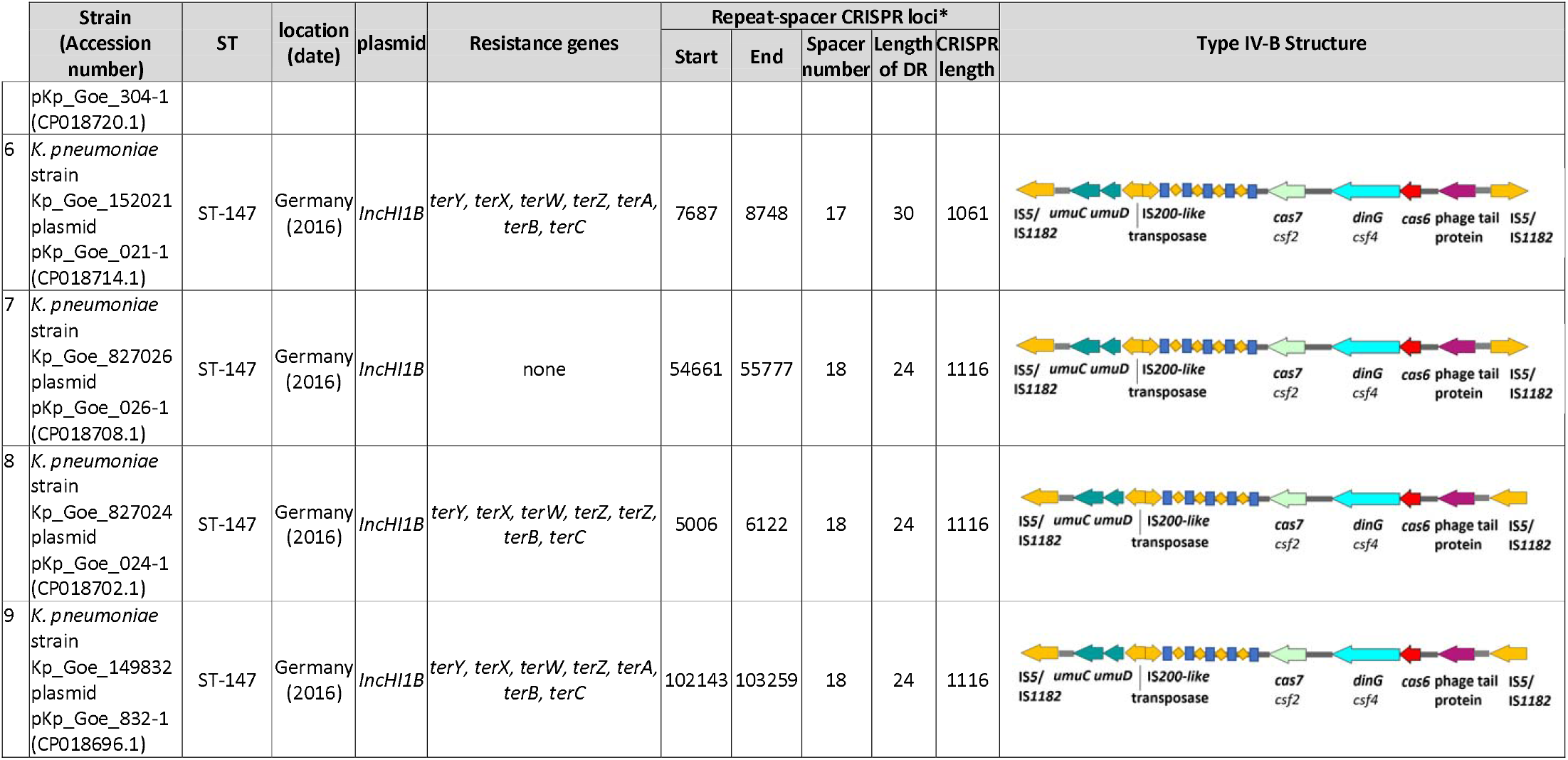
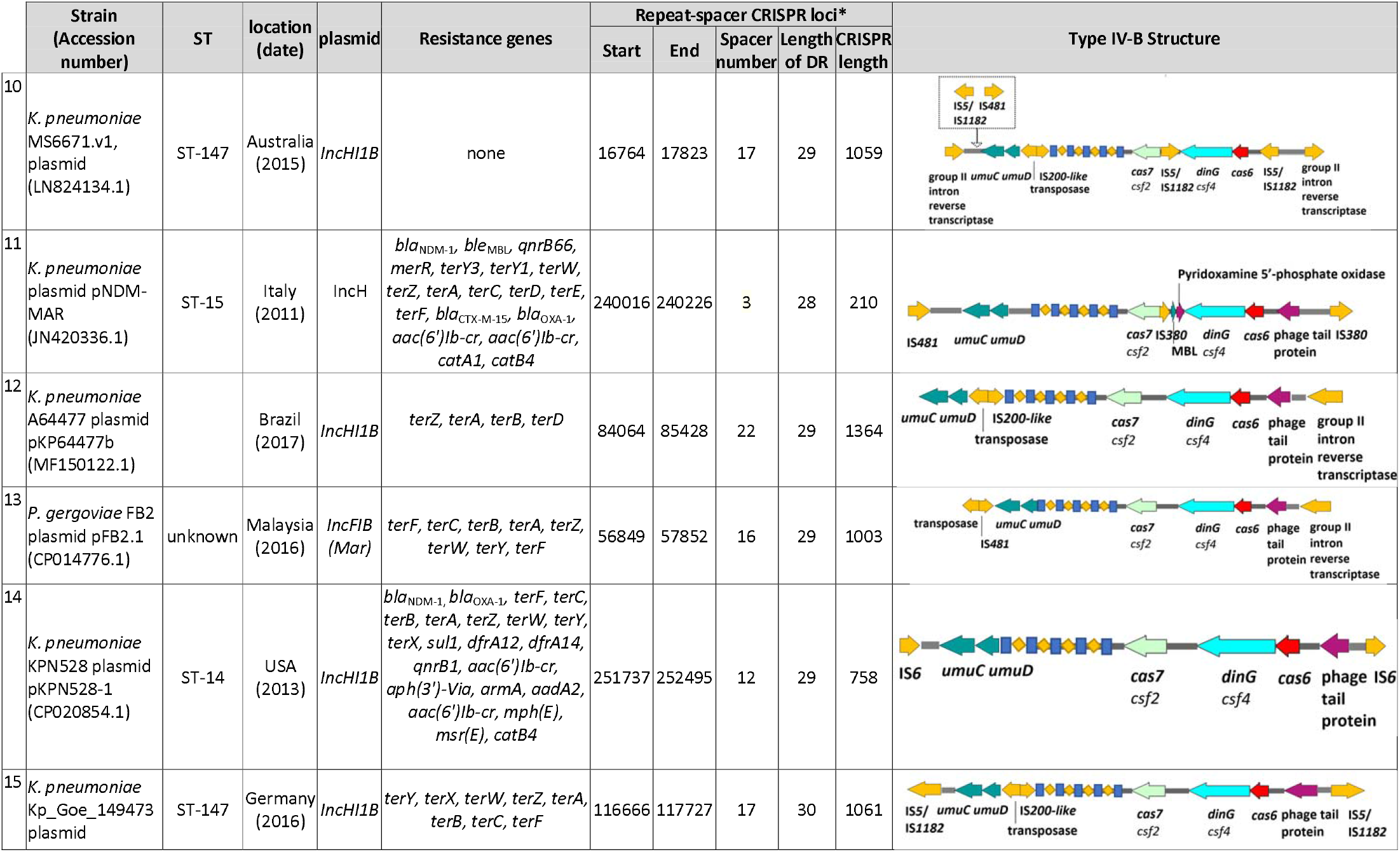
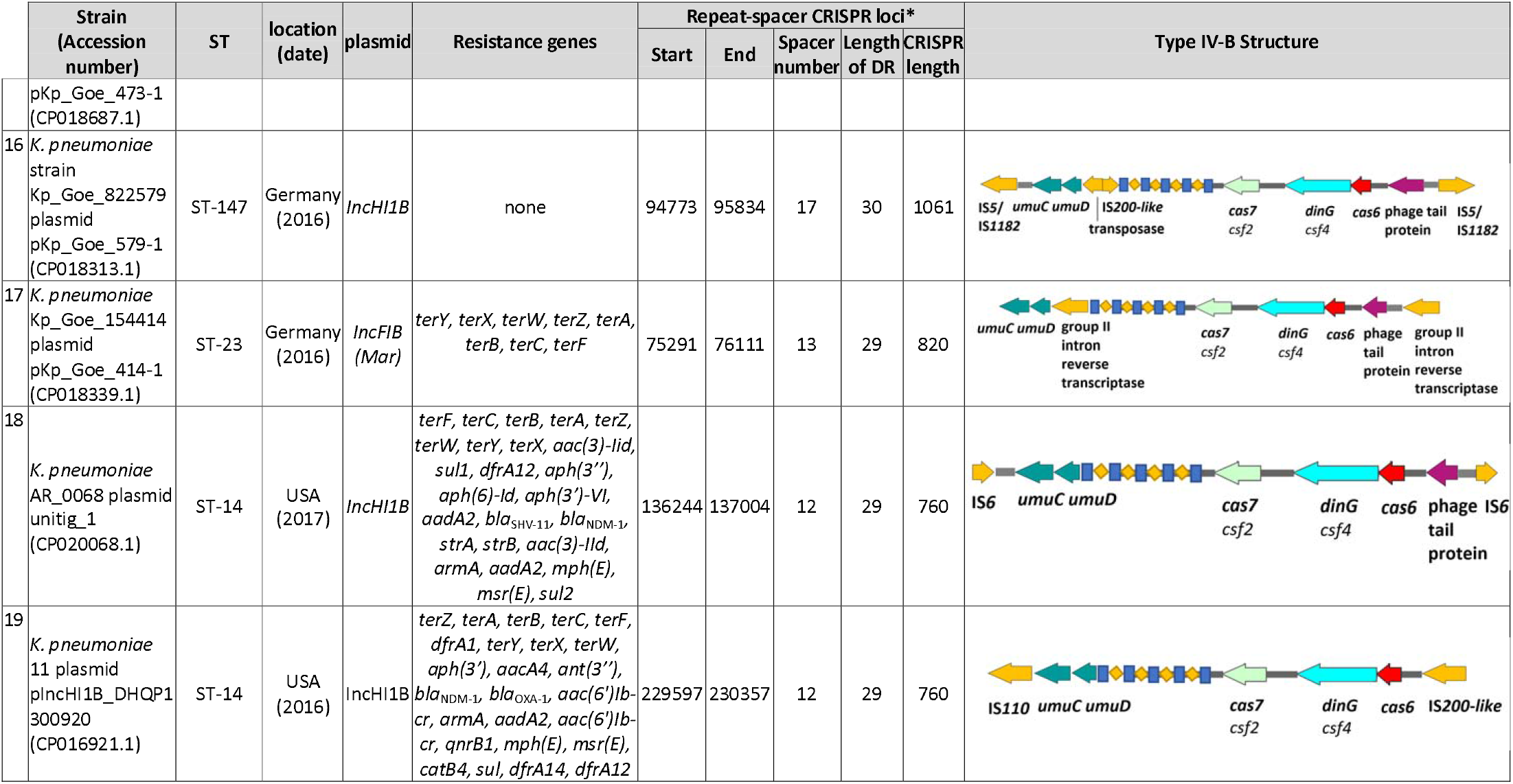
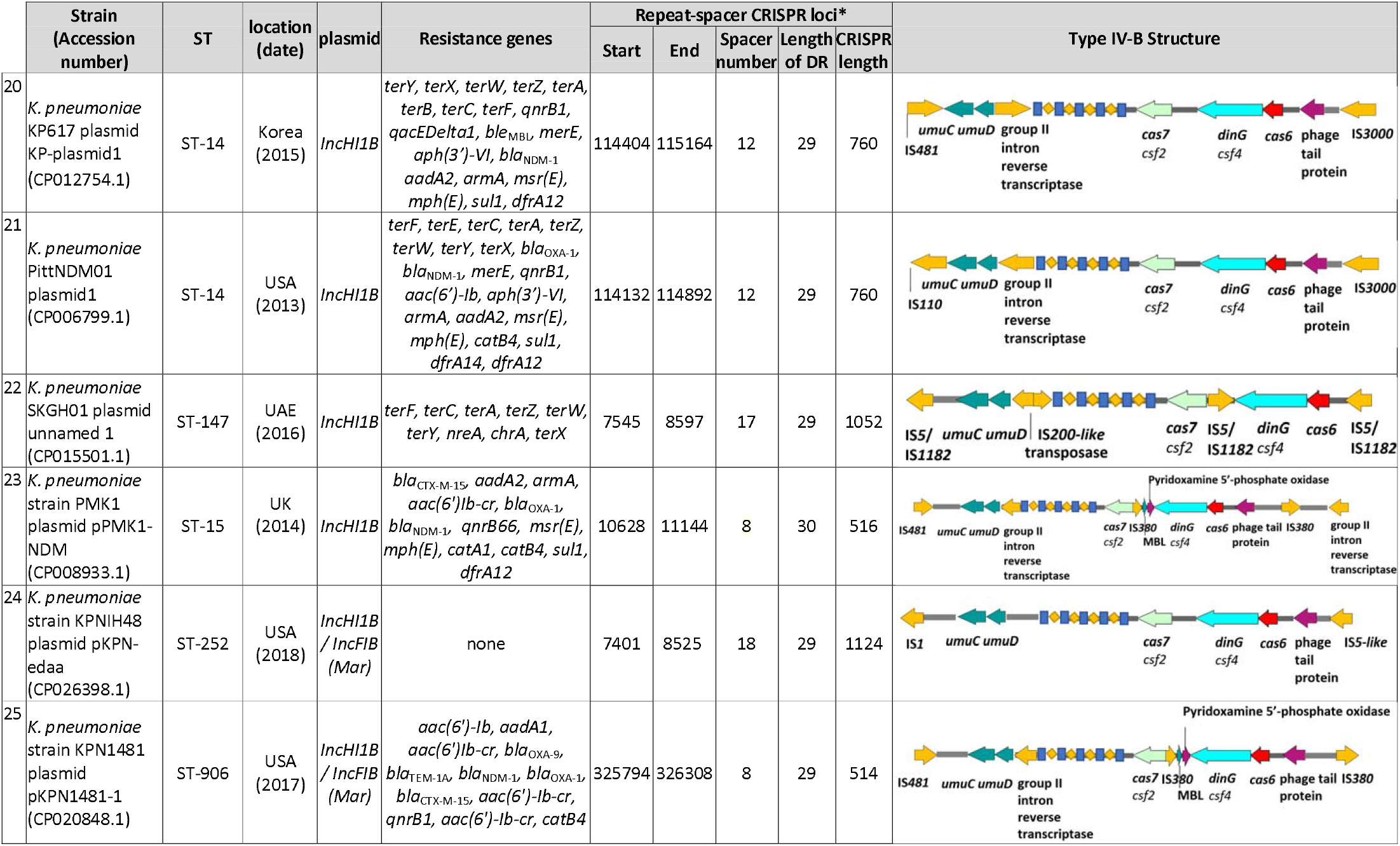
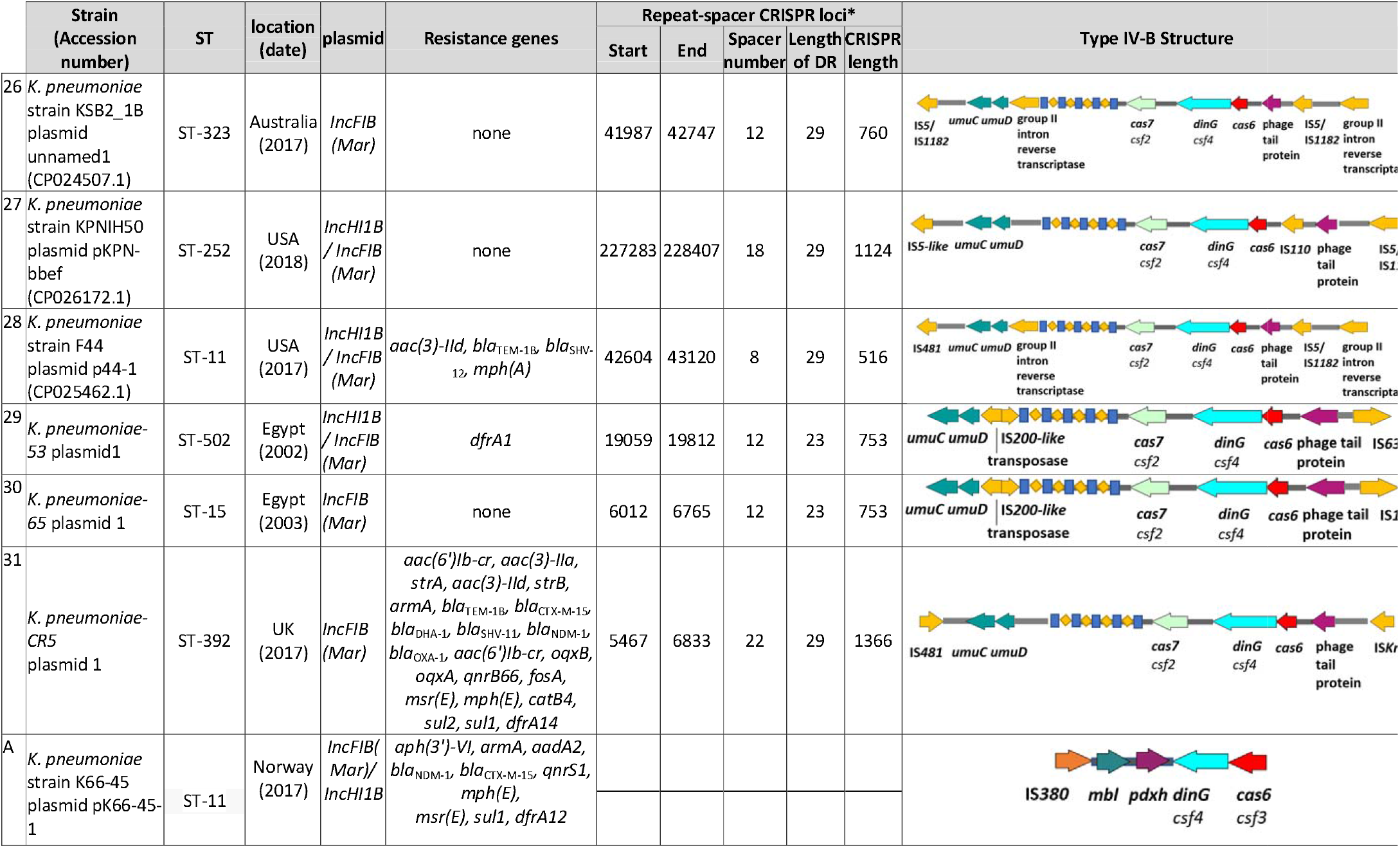
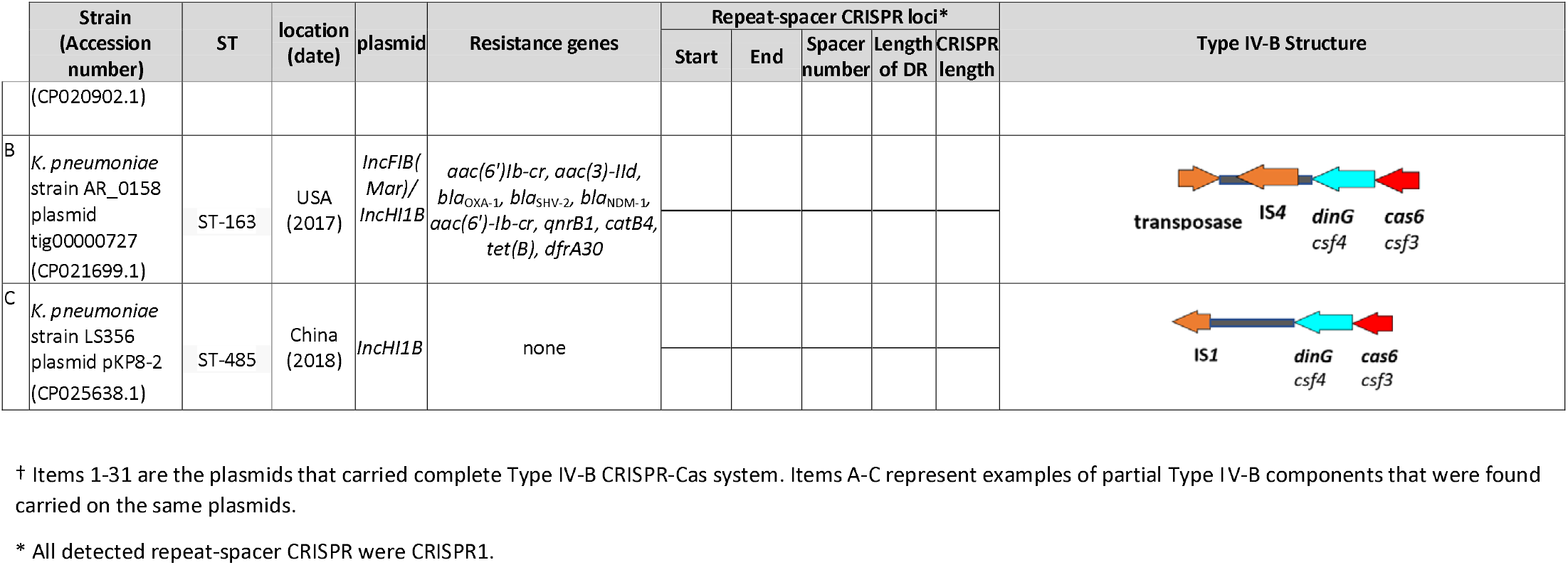
Type IV-B *IncHI1B/IncFIB (Mar)* plasmids information.^†^

